# Spatial Transcriptomics Reveals a Conserved Border Niche and Etiology-Associated Immune Rewiring in Hepatocellular Carcinoma

**DOI:** 10.64898/2026.06.02.729569

**Authors:** Sungwoo Bae, Hongyoon Choi, Su Young Hong, YoungRok Choi, Kwang-Woong Lee, Kwon Joong Na, Suk Kyun Hong

## Abstract

**Background and Aims:** The tumor-stroma interface in hepatocellular carcinoma (HCC) harbors critical intercellular interactions that shape immune evasion and treatment response, yet its spatial architecture remains poorly characterized across etiologies. Whether hepatitis B virus (HBV)-related and non-B non-C (NBNC) HCC share conserved border niche features or exhibit etiology-specific microenvironment programs is unknown. We aimed to spatially resolve the tumor boundary ecosystem and identify etiology-associated signaling networks with translational relevance.

**Approach and Results:** We performed 10x Visium spatial transcriptomics on 11 HCC specimens (7 HBV, 4 NBNC) and applied a machine-learning pipeline integrating CancerFinder and SpaceFlow to define tumor, boundary, and stromal domains. Across etiologies, the boundary zone showed a recurrent desmoplastic niche characterized by cancer-associated fibroblast, tumor-associated macrophage, and tumor endothelial cell accumulation with collagen-integrin extracellular matrix remodeling, including COL1A1-ITGA11 and COL4A1-ITGAV. Etiology-associated differences were observed in the organization of border-zone signaling programs. In representative HBV-related sections, CCL19-CCR7 signaling showed a comparatively restricted, endothelial-skewed topology, whereas representative NBNC sections showed broader inflammatory ligand-receptor networks with elevated NF-kB-associated pathway activity.

**Conclusions:** The HCC tumor-stroma border harbors a recurrent desmoplastic niche upon which etiology-associated immune regulatory programs may be superimposed. These findings generate spatial hypotheses relevant to etiology-informed biomarker development and future therapeutic stratification.

## INTRODUCTION

Hepatocellular carcinoma (HCC) remains a leading cause of cancer-related mortality worldwide, accounting for approximately 750,000 deaths annually and ranking as the third most common cause of cancer death globally [1]. The etiological landscape of HCC is heterogeneous and undergoing a marked epidemiological shift: while chronic hepatitis B virus (HBV) infection continues to drive the majority of cases in East Asia and sub-Saharan Africa, the rising prevalence of metabolic dysfunction-associated steatotic liver disease (MASLD) and alcohol-related liver disease has reshaped the HCC burden in Western populations [2,3]. This etiological diversity carries profound therapeutic implications. The IMbrave150 trial established atezolizumab plus bevacizumab as first-line standard of care for unresectable HCC [4]. However, subsequent analyses have revealed that patients with non-viral etiologies, particularly MASLD-related HCC, derive attenuated benefit from immune checkpoint inhibitor (ICI)-based regimens [5,6]. Corroborating these observations, Pfister et al. demonstrated that non-viral HCC harbors a distinct immunobiological milieu characterized by aberrantly activated CD8+ T cells that paradoxically promote tumor progression [7]. These findings underscore the urgent need to dissect how etiology shapes the tumor microenvironment (TME) at a resolution sufficient to inform precision immunotherapy.

The TME of HCC is among the most complex of solid malignancies, comprising a dynamic interplay of cancer-associated fibroblasts (CAFs), tumor-associated macrophages (TAMs), tumor endothelial cells (TECs), regulatory T cells, and an extensively remodeled extracellular matrix (ECM) [8,9]. Single-cell RNA sequencing (scRNA-seq) studies have substantially advanced our understanding of this complexity, cataloging distinct cell states and identifying key ligand-receptor interactions governing immune evasion in HCC [10,11]. More broadly, single-cell studies across liver cancer and other solid tumors have revealed intratumoral myeloid heterogeneity, myeloid-stromal interactions, exhaustion trajectories of tumor-infiltrating lymphocytes, and candidate immunosuppressive checkpoints beyond PD-1/PD-L1 [12,13]. Nevertheless, a critical limitation pervades these dissociation-based approaches: they obliterate spatial architecture, obscuring where functionally consequential cellular interactions occur. This is particularly significant at the invasive margin, the primary interface of immune-tumor confrontation. Histopathological studies have long recognized that the density and spatial configuration of immune infiltrates at this border zone carry stronger prognostic value than bulk intratumoral assessments [14,15]. Yet the molecular programs governing this niche, and whether they differ by etiology, remain poorly characterized.

Spatial transcriptomics (ST) technologies, exemplified by the 10x Genomics Visium platform, offer a transformative approach by enabling genome-wide expression profiling while preserving tissue architecture [16]. ST has revealed spatially restricted transcriptional programs at tumor boundaries invisible to bulk or single-cell approaches, including tertiary lymphoid structures, hypoxia gradients, and desmoplastic barriers [17,18]. In HCC and primary liver cancer, pioneering ST studies have begun mapping spatial heterogeneity of tumor subclones and the geography of immune exclusion [19,20]. However, existing investigations have predominantly focused on intratumoral heterogeneity or broad tumor-versus-non-tumor comparisons. Prior spatial studies have highlighted immunosuppressive or invasive programs near the HCC margin [19,20], but comparative characterization of the border niche across etiologies -- a key question given reported differences in immunotherapy response -- remains limited. The convergence of machine learning-based spatial domain detection with high-throughput ST now enables objective delineation of the tumor boundary as a discrete analytical unit [21,22].

Here, we present a spatial transcriptomic atlas of 11 surgically resected HCC specimens spanning HBV-associated (n = 7) and non-B non-C (NBNC; n = 4) etiologies, analyzed using 10x Visium coupled with machine learning-driven spatial domain segmentation. By integrating CancerFinder and SpaceFlow algorithms, we objectively define tumor, boundary, and stromal domains and systematically characterize cellular composition, transcriptional programs, and intercellular communication networks [21,22]. Our analysis reveals a conserved desmoplastic border niche -- enriched for CAFs, TAMs, and TECs engaging in collagen-integrin ECM remodeling -- that is shared across etiologies yet is accompanied by etiology-associated differences in immune communication. HBV-associated HCC showed a comparatively narrower, endothelial-skewed border communication topology in representative sections, whereas NBNC-HCC showed broader multicellular inflammatory networks with elevated NF-kB-associated pathway activity.

## Results

### Machine learning-guided spatial domain definition delineates a reproducible tumor-boundary-stroma architecture in hepatocellular carcinoma

To establish an unbiased, spatially resolved framework for dissecting the tumor microenvironment (TME) of hepatocellular carcinoma (HCC), we developed an integrative analytical pipeline combining two complementary machine learning approaches applied to 10x Visium spatial transcriptomics data from 11 resected HCC specimens (7 hepatitis B virus [HBV]-related and 4 non-B non-C [NBNC], including steatosis-associated cases) (**Fig. 1A**). We first applied CancerFinder, a transfer learning-based classifier, to generate spot-level malignancy probability scores across each tissue section. The resulting CancerFinder-derived cancer/normal classification maps reliably distinguished regions of high malignant content from surrounding non-malignant parenchyma in both HBV-related (P1) and NBNC (P10) specimens, as shown alongside the corresponding H&E sections (**Fig. 1B,C**).

**Fig. 1.**
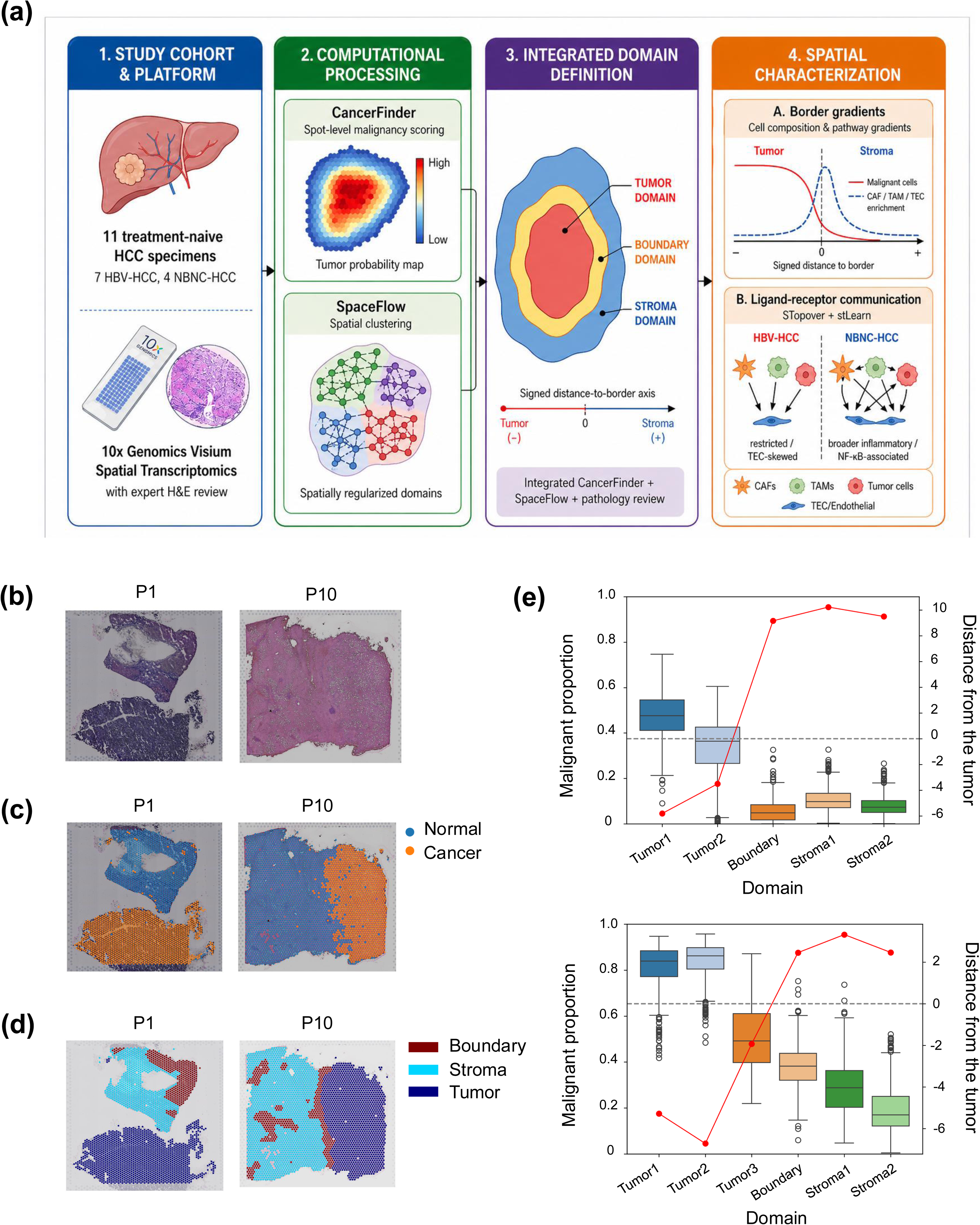
Spatial domain definition and validation in HCC specimens. This figure summarizes the machine learning-guided workflow used to define tumor, boundary, and stromal domains in hepatocellular carcinoma specimens. (**a)** Study workflow. Eleven treatment-naive HCC resection specimens, including seven HBV-related and four non-B non-C cases, were profiled using 10x Visium spatial transcriptomics. CancerFinder was used to estimate spot-level malignancy probability, SpaceFlow was used for spatially regularized clustering, and the resulting outputs were integrated with histologic review to define three spatial domains: Tumor, Boundary, and Stroma. A signed distance-to-border axis was constructed for continuous spatial gradient analyses. (**b)** Hematoxylin and eosin-stained sections from representative HBV-related and NBNC specimens. (**c)** CancerFinder-derived cancer/normal classification maps for the same specimens. Blue indicates normal or low-malignancy-probability regions, and orange indicates cancer or high-malignancy-probability regions. (**d)** Integrated three-domain spatial annotation maps. Dark red indicates Boundary, cyan indicates Stroma, and dark blue indicates Tumor. (**e)** Boxplots of estimated malignant cell proportion across ordered spatial subdomains in the representative specimens. The red line indicates mean distance from the tumor for each subdomain, supporting concordance between the inferred malignancy gradient and the spatial domain hierarchy.

In parallel, we employed SpaceFlow, a spatially regularized deep learning method that integrates gene expression with positional information, to perform unsupervised spatial clustering. To enable systematic cross-sample comparison, we consolidated fine-grained clusters into three principal spatial domains — Tumor, Boundary, and Stroma — guided by the CancerFinder malignancy scores and histological annotation (**Fig. 1D**). The Tumor domain encompassed regions with the highest malignant cell content, the Stroma domain captured the surrounding non-neoplastic liver parenchyma and fibrotic tissue, and the Boundary domain represented the transitional interface between these compartments. Representative spatial domain definitions are shown in **Fig. 1**, and H&E-stained sections for all specimens are provided in **Supplementary Fig. 1**. The malignant-cell gradient across ordered spatial subdomains supported the consistency of the tumor-boundary-stroma framework across all 11 specimens (**Supplementary Fig. 2**).

To validate the biological coherence of the domain assignments, we performed cell-type deconvolution using cell2location with a comprehensive human liver and HCC single-cell RNA sequencing reference atlas [23] [REF]. We then quantified the proportion of malignant cells within each spatial domain and examined its relationship to a signed Euclidean distance-to-border metric, in which negative values denote positions inside the tumor core and positive values denote positions in the peritumoral stroma. As expected, malignant cell proportions were highest in the Tumor domain and declined progressively through the Boundary to the Stroma domain in both representative HBV (P1) and NBNC (P10) specimens (**Fig. 1E**). This gradient was consistent across all 11 samples (**Supplementary Fig. 2**), and the distance-to-border overlay confirmed that the Boundary domain occupied a spatially coherent band at the tumor-stroma interface rather than a diffuse or randomly distributed zone. Taken together, these results establish a robust, ML-guided spatial domain framework that captures the fundamental tissue architecture of HCC and provides a coordinate system for downstream analyses of cellular and molecular gradients across the TME.

### A conserved desmoplastic border niche defined by stromal and myeloid cell enrichment

Having defined three spatial domains, we next asked whether specific cell populations exhibit stereotyped spatial distributions relative to the tumor border, and whether such patterns are conserved across etiologies. Using the cell2location-derived cell-type proportions, we fitted locally weighted scatterplot lowess smoothing curves along the signed distance-to-border axis and compared trends across four representative samples spanning both etiological groups.

Malignant hepatocyte proportions displayed a sharp sigmoidal decline centered on the tumor border, transitioning from high abundance in the tumor core to near-complete absence in the distant stroma in representative specimens (**Fig. 2A**). Cohort-wide malignant cell gradients are summarized in **Supplementary Fig. 2**, and additional distance-to-border cell subtype gradients are shown in **Supplementary Fig. 4**. A reciprocal but less abrupt pattern was observed for tumor-associated T cells (T_tumor), which showed modest enrichment within the tumor domain but gradually diminished with increasing distance into the stroma (**Fig. 2A; Supplementary Figs. 3 and 4**). Critically, this spatial behavior was conserved across all four representative samples irrespective of HBV (P1, P4) or NBNC (P9, P10) status.

**Fig. 2.**
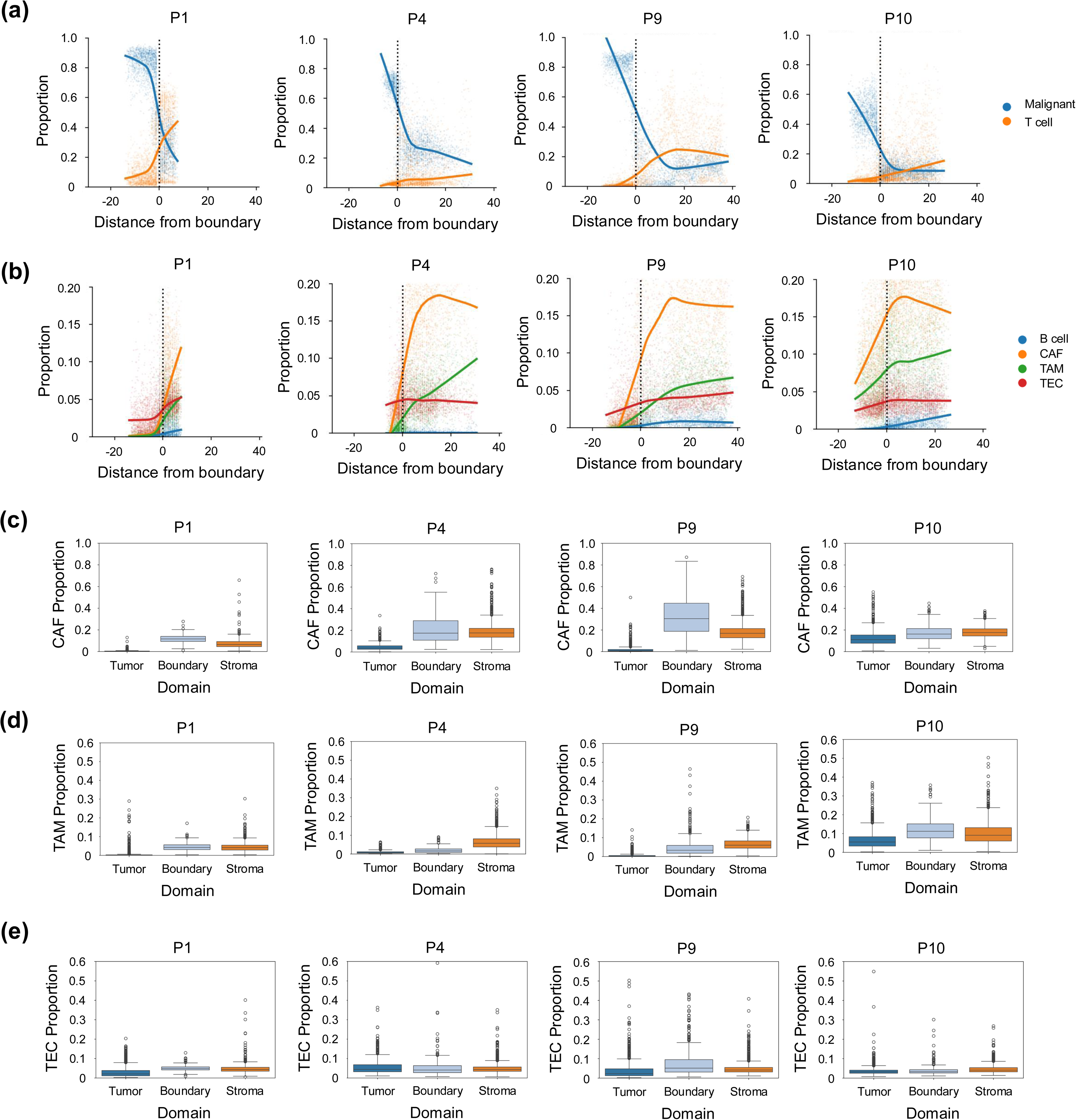
Shared stromal and myeloid remodeling across the tumor-stroma interface. This figure shows cellular gradients and boundary enrichment of stromal and myeloid populations across representative HBV-related and NBNC specimens. **(a)** LOWESS trend curves showing malignant cell and tumor-associated T cell proportions along the signed distance-to-border axis in four representative specimens: P1 and P4 for HBV-related HCC, and P9 and P10 for NBNC-HCC. The vertical dashed line marks the tumor border; negative values indicate tumor-side positions and positive values indicate stromal-side positions. **(b)** LOWESS trend curves for cancer-associated fibroblasts, tumor-associated macrophages, tumor endothelial cells, and tumor-associated B cells along the same distance axis. **(c)** Boxplots comparing cancer-associated fibroblast proportions across Tumor, Boundary, and Stroma domains in the same four representative specimens. **(d)** Boxplots comparing tumor-associated macrophage proportions across the three spatial domains. **(e)** Boxplots comparing tumor endothelial cell proportions across the three spatial domains. The recurrent enrichment of cancer-associated fibroblasts, tumor-associated macrophages, and tumor endothelial cells near the boundary supports a shared stromal-myeloid remodeling program at the HCC tumor-stroma interface.

In contrast to the monotonic gradients of malignant and T_tumor cells, several stromal and myeloid populations exhibited distinctive peaked or enriched distributions centered on or near the Boundary domain. Cancer-associated fibroblasts (CAFs) demonstrated a notable border-centric enrichment pattern, with lowess curves peaking sharply at or immediately adjacent to the tumor-stroma interface and declining in both the tumor-ward and stroma-ward directions across all four representative samples (**Fig. 2B**). Tumor-associated macrophages (TAMs) similarly displayed enrichment in the peritumoral region, though their distribution was broader and extended further into the surrounding stroma compared with CAFs **(Fig. 2B**). Tumor-associated endothelial cells (TECs) also showed preferential accumulation near the boundary, consistent with active neo-angiogenic remodeling at the invasive front (**Fig. 2B**). Tumor-associated B cells (B_tumor) displayed a more variable pattern, with detectable boundary enrichment in some specimens but less consistent peaking across the cohort (**Fig. 2B**).

To quantify these observations, we compared cell-type proportions across the three spatial domains using boxplot analysis. CAF proportions were consistently higher in the Boundary domain than in the adjacent Tumor and Stroma domains across all four representative samples, in keeping with the border-centric pattern observed in the LOWESS analyses (**Fig. 2C**). TAM proportions also showed boundary enrichment, though to a lesser extent in certain NBNC samples (**Fig. 2D**). TEC proportions displayed consistent Boundary enrichment across both etiological groups (**Fig. 2E**). Extended analysis across all 11 specimens showed heterogeneous domain-level distributions of selected lymphoid and endothelial populations, as well as variable distance-to-border gradients for additional cell subtypes (**Supplementary Figs. 3 and 4**).

These findings collectively reveal a conserved desmoplastic border niche in HCC, characterized by the co-accumulation of CAFs, TAMs, and TECs at the tumor-stroma interface. This cellular architecture is reminiscent of the desmoplastic reaction described in other solid malignancies and suggests that the boundary zone functions as a specialized microenvironmental compartment with active stromal remodeling, immune modulation, and angiogenesis, irrespective of HCC etiology.

### Etiology-associated pathway footprints reveal divergent signaling architectures at the tumor border

To characterize the functional signaling landscape across spatial domains, we estimated pathway activity scores for 14 canonical signaling pathways using PROGENy and examined their spatial distributions along the distance-to-border axis. LOWESS trend analysis across four representative samples revealed that several pathways exhibited pronounced spatial gradients with distinct patterns in HBV-related versus NBNC specimens (**Fig. 3A**).

**Fig. 3.**
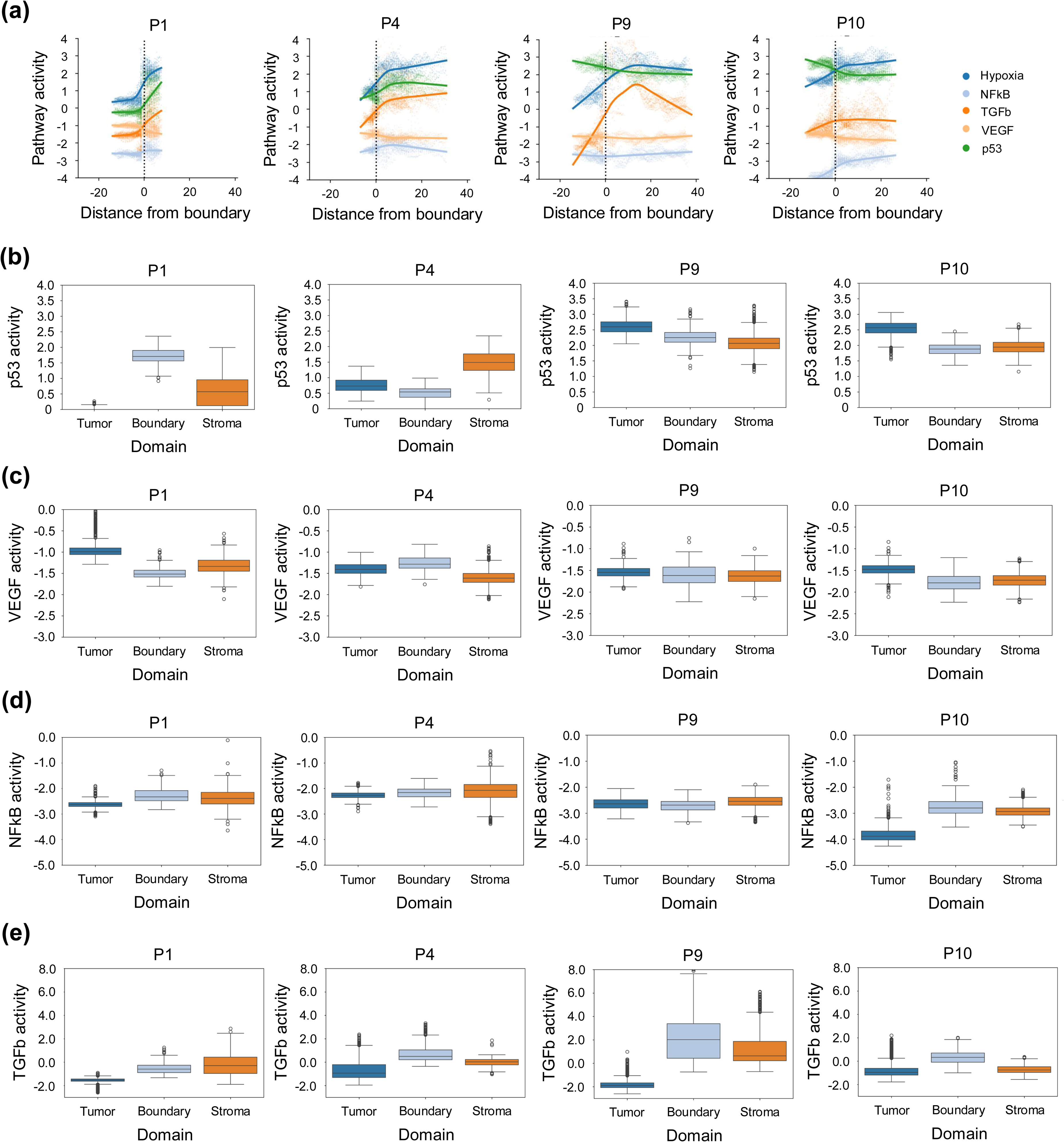
Etiology-associated pathway activity patterns along the tumor-stroma axis. This figure compares pathway activity gradients and domain-level pathway distributions in representative HBV-related and NBNC HCC specimens. (a) LOWESS trend curves showing Hypoxia, NF-kB, TGF-b, VEGF, and p53 pathway activity scores along the signed distance-to-border axis in four representative specimens: P1 and P4 for HBV-related HCC, and P9 and P10 for NBNC-HCC. The vertical dashed line marks the tumor border. (b) Boxplots of p53 pathway activity across Tumor, Boundary, and Stroma domains in the four representative specimens. (c) Boxplots of VEGF pathway activity across the three spatial domains. (d) Boxplots of NF-kB pathway activity across the three spatial domains. (e) Boxplots of TGF-b pathway activity across the three spatial domains. These representative data illustrate spatial heterogeneity in pathway activity across the tumor-stroma axis; cohort-wide domain-level summaries are provided in Supplementary Fig. 5.

Hypoxia pathway activity was consistently elevated within the Tumor domain and declined toward the Stroma in both etiological groups, consistent with the well-characterized hypoxic core of solid tumors (**Fig. 3A**). Similarly, p53 pathway activity displayed a tumor-centric distribution with progressive attenuation across the boundary and into the surrounding stroma, though the magnitude of this gradient varied across specimens. In the four representative specimens, domain-level boxplots showed variable p53 activity patterns across Tumor, Boundary, and Stroma domains, with tumor-side enrichment observed in some but not all specimens (**Fig. 3B**). Cohort-wide domain-level summaries are provided in **Supplementary Fig. 5E**.

VEGF pathway activity, in contrast, exhibited a more nuanced spatial pattern. While generally elevated in the tumor domain, VEGF activity showed a secondary peak or plateau within the Boundary domain in several specimens, particularly in NBNC cases, consistent with active angiogenic signaling at the invasive margin (**Fig. 3A**). In the representative specimens, VEGF activity showed heterogeneous domain-level patterns, with tumor or boundary enrichment in selected cases but without a uniform pattern across all four specimens (**Fig. 3C**). Cohort-wide summaries are provided in **Supplementary Fig. 5C**.

Notable etiology-associated divergence emerged for inflammatory and fibrogenic pathways. NF-kB pathway activity showed divergent spatial profiles between HBV and NBNC groups: HBV-related specimens showed relatively modest NF-kB activity with a flat or gently declining gradient across the tumor-stroma axis, whereas NBNC specimens — particularly those with steatotic backgrounds — exhibited substantially elevated NF-kB activity that was most prominent in the peritumoral Boundary and Stroma domains (**Fig. 3A**). In the representative specimens, NF-kB activity showed more prominent peritumoral or stromal enrichment in NBNC sections, particularly P10, than in the HBV-related sections shown (**Fig. 3D**). This representative pattern should be interpreted together with the cohort-wide summaries in **Supplementary Fig. 5B**. This finding aligns with the established role of NF-kB-mediated chronic inflammation in steatosis-driven hepatocarcinogenesis.

TGF-b pathway activity also showed spatial heterogeneity across the tumor-stroma axis. In the representative HBV-related specimens, TGF-b activity was relatively uniform or mildly enriched near the Boundary domain, whereas selected NBNC specimens showed more prominent peritumoral enrichment (**Fig. 3A and 3E**). These observations suggest a fibrogenic overlay at the border in some NBNC tumors but should be interpreted cautiously given the limited number and heterogeneity of NBNC cases. Additional pathway summaries across the complete cohort are provided in **Supplementary Fig. 5**, including exploratory JAK-STAT pathway activity distributions (**Supplementary Fig. 5F**).

Together, these pathway-level analyses support the interpretation that while certain tumor-intrinsic programs (hypoxia, p53) are spatially conserved across etiologies, the inflammatory and fibrogenic signaling architecture of the HCC border zone differs between HBV-related and NBNC tumors. The heightened NF-kB and TGF-b activity in NBNC specimens is consistent with a more pronounced inflammatory and fibrogenic overlay at the border in non-viral HCC.

### Border-enriched ligand-receptor interactions reveal a conserved ECM scaffold with etiology-associated immune communication networks

To investigate the intercellular communication programs operating within the desmoplastic border niche, we performed ligand-receptor (L-R) interaction analysis using a multi-method framework comprising LIANA+ for interaction prioritization, stLearn for spatially resolved cell-type network inference, and STopover [24] for topological overlap assessment of ligand and receptor expression domains. We ranked L-R pairs by their STopover Jaccard composite scores within the Boundary domain, designating interactions with scores exceeding 0.3 as strongly border-enriched and those with lower but spatially concentrated overlap as boundary-associated.

Functional enrichment analysis of border-enriched L-R interactions in a representative specimen (P4) revealed a dominant overrepresentation of terms related to extracellular matrix (ECM) organization, collagen biosynthesis, integrin-mediated signaling, and cell-matrix adhesion (**Fig. 4A**). This ECM-centric signature was shared across both HBV and NBNC specimens, identifying the collagen-integrin signaling axis as a conserved molecular feature of the HCC border niche. Among the prominent boundary-associated interactions, COL1A1-ITGA11 showed spatially concentrated ligand-receptor overlap at the tumor-stroma interface, with STopover mapping revealing focal overlap within the Boundary domain despite a Jaccard composite score below the predefined strong-enrichment threshold (**Fig. 4B**). The boundary-associated COL4A1-ITGAV interaction similarly exhibited spatially concentrated overlap near the tumor-stroma interface, suggesting that fibrillar and basement membrane collagen-integrin interactions are recurrent components of the invasive-margin niche (**Fig. 4C**). Spatially resolved cell-type network analysis using stLearn for the COL1A1-ITGA11 interaction revealed a putative communication network involving stromal, malignant, fibroblast, epithelial, and endothelial compartments (**Fig. 4D**). This network architecture suggests that stromal collagen-integrin signaling may coordinate ECM remodeling and endothelial or fibroblast-associated programs within the border niche.

**Fig. 4.**
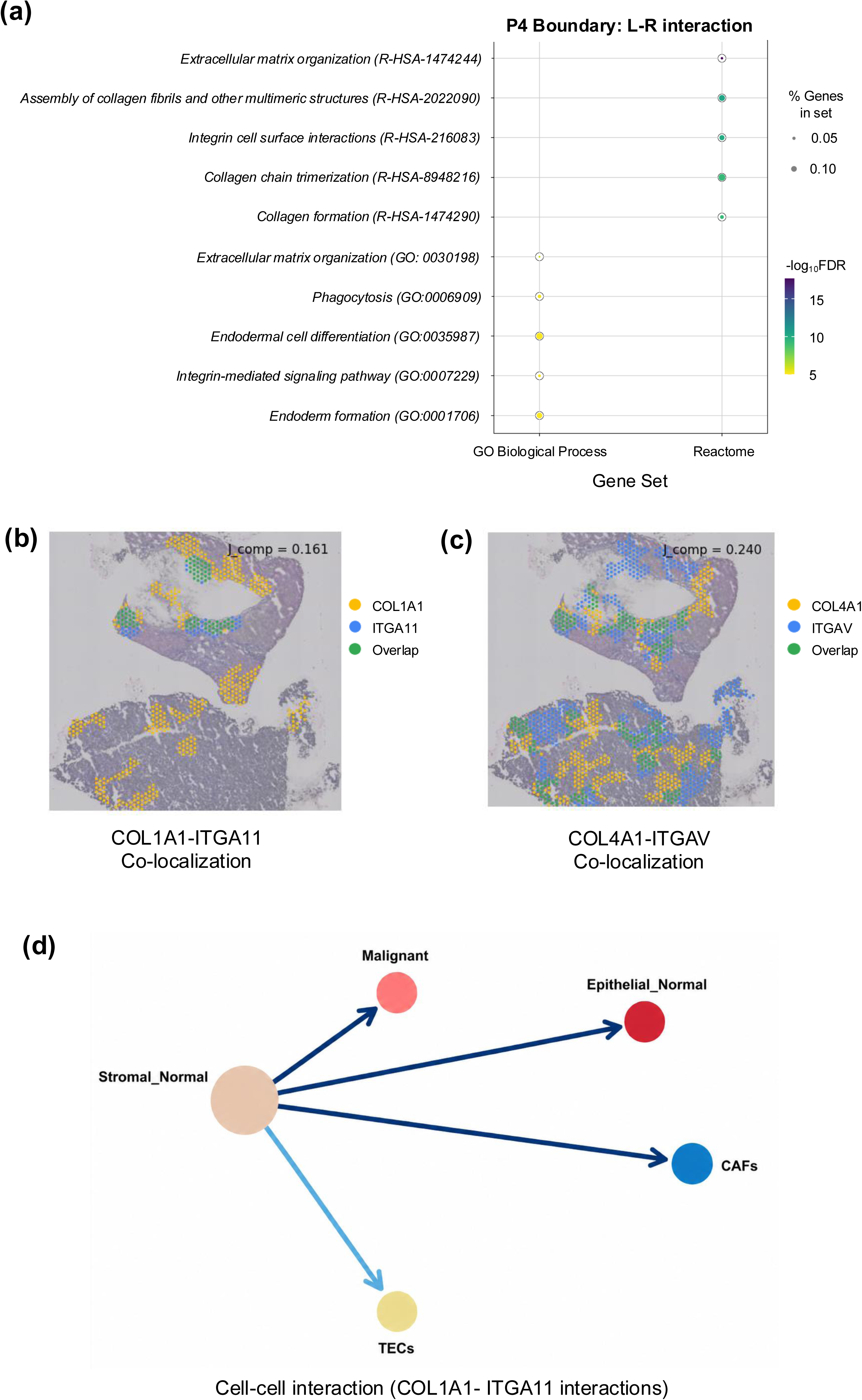
Conserved extracellular matrix remodeling at the HCC tumor boundary. This figure highlights collagen-integrin ligand-receptor interactions as a recurrent component of the HCC boundary niche. **(a)** Dot plot of functional enrichment terms for boundary-enriched ligand-receptor interactions in representative specimen P4. Dot size indicates the percentage of genes represented in each gene set, and color indicates the negative log-transformed false discovery rate. **(b)** STopover spatial colocalization map for the COL1A1-ITGA11 interaction in P4. Yellow/orange spots indicate ligand expression, blue spots indicate receptor expression, and green spots indicate spatial overlap between ligand and receptor expression domains. The Jaccard composite score is shown on the map. **(c)** STopover spatial colocalization map for the COL4A1-ITGAV interaction, displayed with the same color scheme. **(d)** stLearn-inferred cell-cell interaction network for the COL1A1-ITGA11 axis. Nodes indicate inferred cell populations, and arrows indicate predicted directional ligand-receptor communication; arrow thickness reflects relative interaction strength. These results support collagen-integrin signaling as a boundary-associated communication axis involved in extracellular matrix remodeling at the HCC tumor-stroma interface. Raw cell-state labels in the network correspond to the cell2location reference atlas: Stromal_Normal, non-tumor stromal cells; Epithelial_Normal, non-tumor epithelial cells; CAFs, cancer-associated fibroblasts; TECs, tumor endothelial cells.

While the ECM scaffold was conserved, the immune communication networks at the tumor border diverged between etiological groups. Functional enrichment analysis of border-enriched L-R interactions in a representative HBV specimen (P6) identified terms predominantly related to ECM organization, with additional representation of immune system processes and signal transduction pathways (**Fig. 5A**). In contrast, a representative NBNC specimen (P10) displayed a distinct functional profile, with pronounced enrichment of cytokine signaling, inflammatory response, and immune system regulation terms among its border-enriched interactions (**Fig. 5B**). This distinction implies that the molecular vocabulary of the border niche shifts from structurally dominated communication in HBV-related tumors toward a more immunologically active signaling repertoire in NBNC tumors.

**Fig. 5.**
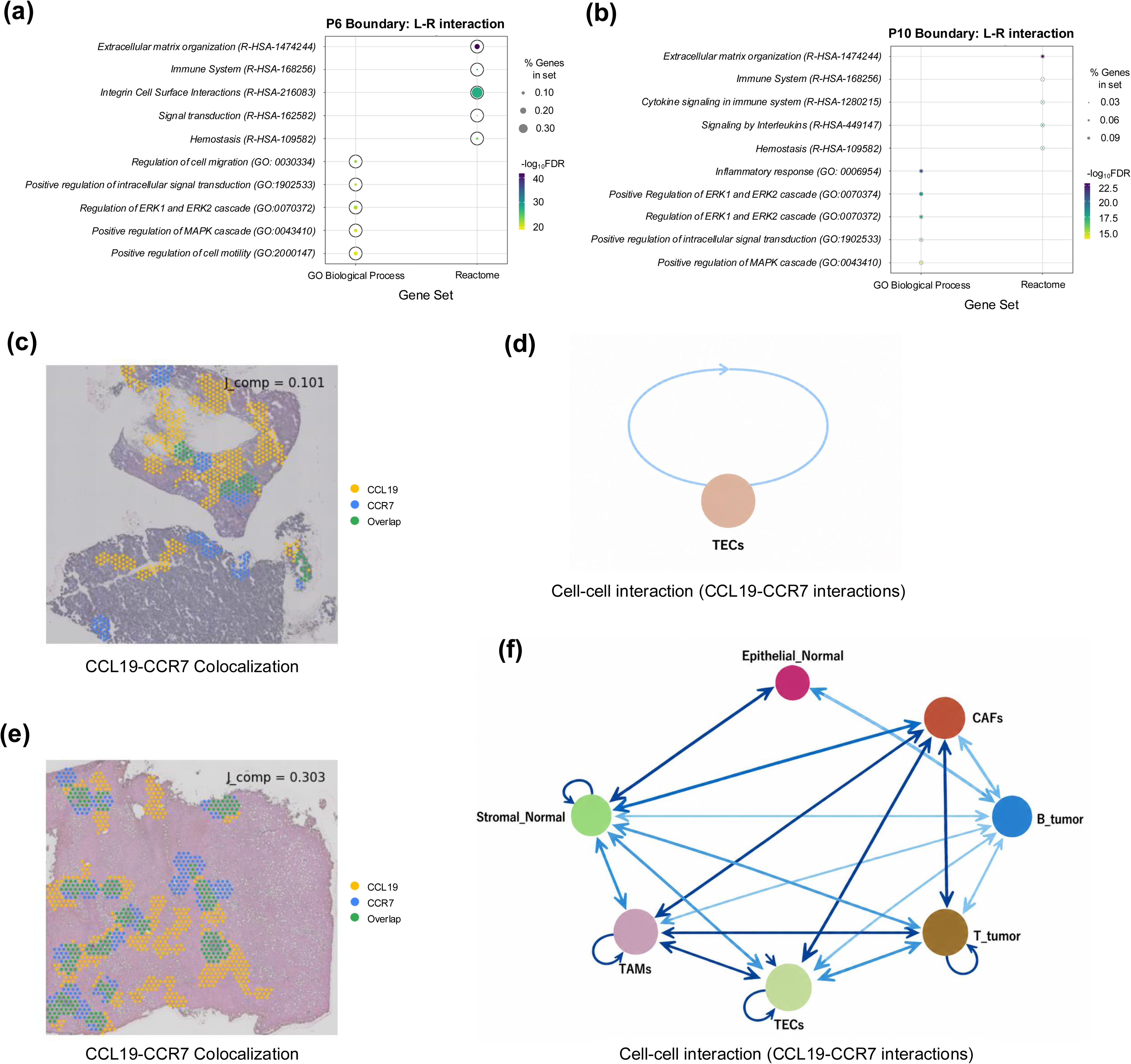
Etiology-associated immune communication networks at the HCC tumor boundary. This figure contrasts ligand-receptor enrichment and CCL19-CCR7 communication topology between representative HBV-related and NBNC HCC specimens. (a) Dot plot of functional enrichment terms for boundary-enriched ligand-receptor interactions in HBV-related specimen P6. Dot size indicates the percentage of genes represented in each gene set, and color indicates the negative log-transformed false discovery rate. (b) Dot plot of functional enrichment terms for boundary-enriched ligand-receptor interactions in NBNC specimen P10, showing enrichment of immune and inflammatory signaling terms. (c) STopover spatial colocalization map for CCL19-CCR7 in HBV-related P6. Yellow/orange spots indicate CCL19 expression, blue spots indicate CCR7 expression, and green spots indicate ligand-receptor spatial overlap. The Jaccard composite score is shown on the map. (d) stLearn-inferred CCL19-CCR7 interaction network in HBV-related P6, showing a restricted topology centered on tumor endothelial cell-associated signaling. € STopover spatial colocalization map for CCL19-CCR7 in NBNC P10, displayed with the same color scheme. (f) stLearn-inferred CCL19-CCR7 interaction network in NBNC P10, showing broader multicellular communication involving stromal, fibroblast, macrophage, endothelial, epithelial, and tumor-associated lymphoid compartments. Raw cell-state labels in the network correspond to the cell2location reference atlas: B_tumor, tumor-associated B cells; T_tumor, tumor-associated T cells; Stromal_Normal, non-tumor stromal cells; and Epithelial_Normal, non-tumor epithelial cells. The contrast between P6 and P10 supports etiology-associated differences in the spatial organization of chemokine signaling at the HCC tumor boundary.

To illustrate this divergence at the level of individual L-R pairs, we examined the CCL19-CCR7 axis, a chemokine interaction critically involved in lymphocyte trafficking and tertiary lymphoid structure organization. In the HBV specimen (P6), STopover analysis of CCL19-CCR7 yielded a relatively low Jaccard composite score (J_comp = 0.101), with spatially restricted overlap confined to small foci within the boundary zone (**Fig. 5C**). The corresponding stLearn network revealed a limited communication architecture centered on a TEC autocrine loop with sparse connections to other cell populations (**Fig. 5D**). In this representative HBV section, CCL19-CCR7 displayed a comparatively narrow, endothelial-skewed topology with limited capacity for broad immune cell recruitment.

In contrast, the same CCL19-CCR7 interaction in the NBNC specimen (P10) achieved a substantially higher topological overlap score (J_comp = 0.303), with STopover mapping revealing broadly distributed spatial colocalization across the Boundary domain and extending into the adjacent stroma (**Fig. 5E**). The stLearn network for this specimen demonstrated a broader multicellular communication topology, with CCL19-CCR7 signaling involving stromal, fibroblast, macrophage, endothelial, epithelial, and tumor-associated lymphoid compartments in a densely interconnected network (**Fig. 5F**). This architecture suggests that NBNC-associated HCC may support a broader CCL19-CCR7 signaling circuit at the tumor border, potentially facilitating enhanced immune cell recruitment, organization, and crosstalk within the desmoplastic niche.

The juxtaposition of these findings supports a dual-layered organization of intercellular communication at the HCC border: a conserved ECM-integrin scaffold that defines the structural framework of the desmoplastic niche across etiologies, overlaid with etiology-associated immune signaling differences. HBV-related tumors exhibit a comparatively narrower border characterized by restricted chemokine signaling and narrow cell-type engagement, whereas NBNC tumors — particularly those arising in steatotic livers — deploy expansive inflammatory and chemokine communication networks that recruit diverse cellular participants into the border niche. These etiology-associated immune circuits may be relevant to reported heterogeneity in immunotherapy responses and represent candidate targets for future etiology-stratified investigation in HCC.

## Discussion

The tumor border represents a critical interface where malignant expansion confronts host tissue defense, yet the molecular architecture of this niche in hepatocellular carcinoma (HCC) remains incompletely characterized, particularly with respect to etiological context. In this study, we leveraged spatial transcriptomics across 11 HCC specimens spanning HBV-related and non-B non-C (NBNC) etiologies to delineate the ligand-receptor networks and pathway activities that define the tumor-stroma boundary. Our analyses support a two-tiered organization of the HCC border niche: a conserved desmoplastic scaffold characterized by CAF, TAM, and TEC accumulation with collagen-integrin extracellular matrix remodeling, overlaid by etiology-associated immune communication states. Within this framework, HBV-associated tumors suggested a comparatively narrower, endothelial-skewed communication topology, whereas NBNC tumors exhibited broader inflammatory and fibrogenic rewiring. These findings provide a spatially resolved framework for generating testable hypotheses about how etiology may shape the immune-regulatory architecture of the tumor border.

The identification of a conserved desmoplastic border niche across all 11 specimens aligns with the broader concept that solid tumors co-opt wound-healing and matrix-remodeling programs at their invasive front [25,26]. In our data, the most reproducible shared feature was the boundary-associated enrichment of collagen-integrin interactions (COL1A1-ITGA11, COL4A1-ITGAV, COL1A2-ITGAV) together with CAF, TAM, and TEC accumulation. This is consistent with prior evidence that stromal α11β1 is a fibrillar collagen receptor enriched in CAFs and contributes to collagen reorganization, stromal stiffness, and tumor progression [27], and with the paradigm that integrin αV heterodimers link matrix stiffness to TGF-b activation [28]. TGF-b activity was also detected at the border, but its prominence appeared more variable across etiologies and was particularly accentuated in NBNC specimens. Accordingly, the shared border program may be interpreted most conservatively as a collagen-remodeling desmoplastic scaffold, with TGF-b representing a context-dependent fibrogenic reinforcement rather than the defining universal driver [29,30]. The co-localization of Kupffer-like TAMs within this niche is notable, as tissue-resident macrophage populations in the liver possess unique tolerogenic properties that may be subverted at the tumor border [31]. From a therapeutic perspective, the conserved collagen-integrin/TGF-b remodeling axis warrants further investigation as a candidate etiology-agnostic stromal target, although therapeutic translation will require prospective validation [32,33].

Superimposed on this shared scaffold, HBV-associated tumors suggested a comparatively narrower, endothelial-skewed border communication topology. Although p53 and VEGF displayed spatial gradients across the tumor-stroma axis, these patterns were primarily domain-associated rather than consistently etiology-specific across the full cohort and should therefore be interpreted cautiously [34,35]. In representative HBV-related sections, the CCL19-CCR7 axis showed a TEC-centered configuration with relatively limited multicellular spread, consistent with a more spatially restricted communication state at the border. CCL19-CCR7 signaling is canonically associated with lymphocyte homing and has been linked to favorable prognosis in several cancers [36]; however, its inferred autocrine engagement within endothelial cells, rather than in a paracrine lymphocyte-recruiting configuration, may reflect a co-opted function potentially relevant to vascular permeability or immune cell transit [37]. Additional inferred immunoregulatory interactions, including LGALS9- and CD47-related signals, were also observed in some HBV sections, but these will require orthogonal validation before stronger mechanistic conclusions can be drawn [38,39]. More broadly, this endothelial-skewed border state should be interpreted in light of the established coupling between tumor angiogenesis, vascular remodeling, and immunosuppression in the tumor microenvironment [40].

In contrast, NBNC tumors in our cohort, several of which arose in steatotic liver backgrounds, displayed a broader inflammatory and fibrogenic border state characterized by higher NF-kB activity and a more densely interconnected multicellular communication topology. These findings are compatible with prior reports linking steatotic or non-viral HCC to inflammatory immune dysregulation [41], but direct equivalence with MASLD- or NASH-HCC should be avoided in this cohort. Additional inferred chemotactic and innate immune-associated interactions, including CXCL12-CXCR4 and BGN-TLR2, were present in some sections [42,43], supporting the interpretation that NBNC border biology may involve broader inflammatory recruitment and signaling. This is consistent with the observation by Pfister et al. that NASH-related HCC harbors aberrantly activated, exhausted CD8+ T cells and that patients with non-viral HCC may derive reduced benefit from immune checkpoint inhibitor monotherapy [7]. Our data add a spatial layer by suggesting that inflammatory signaling in NBNC-HCC may be concentrated at the border through specific ligand-receptor circuits involving CAFs, TAMs, TECs, and epithelial cells.

These etiology-associated border states may have translational implications for combination immunotherapy design in HCC. Because this cohort consisted of treatment-naive resected tumors without matched outcome data, our study does not establish treatment-response mechanisms directly. Instead, the data provide a spatial framework that may help contextualize reported heterogeneity in immunotherapy outcomes across etiologies [4,6]. In this framework, HBV-HCC may involve a relatively more endothelial-centered border state, whereas NBNC-HCC may be shaped by broader inflammatory and chemotactic rewiring. These border-state differences generate testable hypotheses for future biomarker-stratified studies.

Several limitations of this study warrant consideration. First, our cohort of 11 specimens, while sufficient for a spatial transcriptomics discovery study given the technical complexity and cost of the platform, limits statistical power for detecting more subtle etiology-associated differences and precludes formal subgroup analyses within the NBNC category, which itself encompasses heterogeneous metabolic, cryptogenic, and potentially alcohol-related etiologies. NBNC encompasses heterogeneous backgrounds including but not limited to steatotic liver disease; direct equivalence with MASLD or NASH should not be assumed. Nevertheless, the repeated observation of boundary-associated stromal and myeloid enrichment across specimens supports the robustness of the core spatial pattern, whereas etiology-associated differences should be interpreted as hypothesis-generating. Second, our findings await orthogonal validation by complementary spatial methods such as multiplex immunohistochemistry or CODEX imaging, which would provide protein-level and single-cell resolution confirmation of the ligand-receptor co-localization patterns inferred from transcriptomic data. Furthermore, ligand-receptor interactions inferred from spot-level spatial transcriptomics represent co-localization patterns rather than confirmed functional signaling events. Third, the absence of matched clinical outcome data in this cohort prevents direct correlation of border niche features with treatment response, and prospective studies linking spatial border phenotypes to immunotherapy outcomes are needed to establish clinical utility. Because the cohort consists of treatment-naive resected tumors, direct extrapolation to systemic therapy response in advanced unresectable HCC should be made cautiously. Fourth, the characterization of NBNC-HCC with steatotic background is an area of active clinical urgency as this etiology rises in global prevalence [44], and even preliminary spatial molecular characterization such as that provided here can inform hypothesis generation for the design of dedicated trials. Rather than directly proving differential therapy response, this study provides a spatial framework for understanding how etiology may shape the immune-regulatory architecture of the HCC border niche, generating testable hypotheses for etiology-informed therapeutic stratification.

## Methods

### Study Population and Specimen Collection

This study enrolled treatment-naive patients undergoing curative-intent hepatic resection for hepatocellular carcinoma (HCC) at Seoul National University Hospital. The study protocol was approved by the Institutional Review Board, and written informed consent was obtained from all participants. A total of eleven HCC specimens were included, comprising seven hepatitis B virus (HBV)-related cases and four non-B non-C (NBNC) cases with underlying hepatic steatosis. Specimens are identified by sample identifiers P1 through P11. Patients who had received prior locoregional or systemic therapy were excluded.

### Tissue Processing and Spatial Transcriptomics

Fresh surgical specimens were procured immediately following resection. Tissue blocks (∼5 mm3) were excised from the tumor-normal interface to capture the spatial continuum from tumor core through boundary zone into non-neoplastic parenchyma. Specimens were OCT-embedded, snap-frozen, and cryosectioned at 10 um onto Visium Spatial Gene Expression slides (10x Genomics). Spatially barcoded gene expression profiling was performed using the 10x Visium platform, with libraries sequenced on an Illumina NovaSeq system. Raw data were processed using Space Ranger v2.0.0 with alignment to GRCh38, and quality control filtering removed spots with low UMI counts, few detected genes, or elevated mitochondrial transcript proportions.

### Spatial Domain Annotation

To define biologically meaningful spatial domains, we employed a two-pronged strategy. CancerFinder [21], a transfer learning framework for tumor cell annotation, assigned a malignancy probability score to each Visium spot. SpaceFlow [22], an unsupervised spatial clustering method based on graph convolutional networks, identified spatially coherent transcriptomic domains. Integrating both outputs, spots with malignancy probability exceeding 50% and spatial continuity with histologically reviewed tumor regions were classified as the Tumor domain. SpaceFlow-derived clusters directly adjacent to the tumor domain and located at the histologically reviewed tumor-stroma interface were assigned to the Boundary domain, particularly when they showed intermediate malignancy probability or spatial continuity with both tumor and stromal compartments. Remaining non-neoplastic tissue was classified as the Stroma domain. A signed Euclidean distance-to-border metric was computed for each spot to enable continuous spatial analyses.

### Cell-Type Deconvolution

Cell-type composition within each Visium spot was estimated using cell2location, a Bayesian deconvolution model integrating single-cell RNA sequencing reference data with spatial transcriptomics. The reference dataset comprised curated scRNA-seq profiles from human liver and HCC tissues, encompassing immune, stromal, and parenchymal populations.

### Pathway Activity and Spatial Modeling

Pathway-level signaling activity was quantified using PROGENy for 14 canonical signaling pathways. Cell-type contributions to pathway activity were additionally explored using MISTy.

### Ligand-Receptor Interaction Analysis

Spatially informed cell-cell communication was assessed through a multi-method framework. LIANA+ provided consensus rankings of ligand-receptor pairs by integrating multiple interaction databases and scoring methods. stLearn incorporated spatial proximity constraints to prioritize interactions between physically co-localized cell populations. STopover applied topological spatial overlap analysis; interactions were ranked by Jaccard composite score within the Boundary domain; those exceeding 0.3 were designated strongly border-enriched, while lower-scoring interactions with spatially concentrated boundary overlap were classified as boundary-associated.

### Statistical Analysis

Given the exploratory nature of this study and the modest sample size (n = 11), the statistical framework prioritized descriptive characterization, effect size estimation, and concordance across analytical modalities. Comparisons across spatial domains and distance-to-border bins were performed using the Wilcoxon rank-sum test and the Kruskal-Wallis test. Trends were assessed for concordance across multiple independent analytical approaches to reinforce biological conclusions through methodological triangulation. Detailed descriptions of all computational methods are provided in **Supplementary Methods**.

### Statement of Ethics

This study was approved by the Institutional Review Board of Seoul National University Hospital, Seoul, Republic of Korea (IRB No. H-2204-047-1314). Written informed consent was obtained from all participants.

## Data Availability Statement

Processed Visium AnnData objects of this cohort and clinical metadata necessary to are available to qualified investigators on request, subject to institutional data-sharing policies.

## Author Contributions (CRediT)

Conceptualization: Sungwoo Bae, Suk Kyun Hong, Kwon Joong Na

Methodology: Sungwoo Bae, Hongyoon Choi, Kwon Joong Na

Software: Sungwoo Bae, Hongyoon Choi

Validation: Su Young Hong, YoungRok Choi, Kwang-Woong Lee, Suk Kyun Hong

Formal Analysis: Sungwoo Bae, Kwon Joong Na

Investigation: Sungwoo Bae, Hongyoon Choi, Kwon Joong Na, Suk Kyun Hong

Resources: Su Young Hong, YoungRok Choi, Kwang-Woong Lee, Suk Kyun Hong

Data Curation: Su Young Hong, YoungRok Choi, Kwang-Woong Lee, Suk Kyun Hong, Kwon Joong Na

Writing - Original Draft: Sungwoo Bae, Kwon Joong Na

Writing - Review & Editing: Sungwoo Bae, Hongyoon Choi, Su Young Hong, YoungRok Choi, Kwang-Woong Lee, Suk Kyun Hong, Kwon Joong Na

Visualization: Sungwoo Bae, Kwon Joong Na

Supervision: Su Young Hong, YoungRok Choi, Kwang-Woong Lee, Suk Kyun Hong

Project Administration: Suk Kyun Hong, Kwon Joong Na

Funding Acquisition: Hongyoon Choi, Kwon Joong Na, Suk Kyun Hong

## AI/LLM disclosure

During manuscript preparation, a generative AI tool was used for language editing. The authors reviewed, edited, and verified all content and take full responsibility for the final manuscript.

