## Supplementary Data for "Spatial Transcriptomics Reveals a Conserved Border Niche and Etiology-Associated Immune Rewiring in Hepatocellular Carcinoma"

**Supplementary Materials**

**Supplementary Table**

**Supplementary Figures**

**Supplementary References**

**Supplementary Methods**

***CancerFinder: Tumor Cell Classification***

CancerFinder is a transfer learning framework designed for tumor cell annotation in transcriptomic data [S1]. The method leverages pre-trained models derived from large-scale cancer transcriptomic datasets (including TCGA and GEO repositories) to discriminate malignant from non-malignant expression signatures. For each Visium spot, CancerFinder generates a continuous probability estimate of neoplastic identity by projecting the spot-level expression profile into a learned latent space trained on thousands of tumor and normal samples. The malignancy probability threshold of 50% was selected based on the bimodal distribution of scores observed across specimens, which consistently separated histologically confirmed tumor and non-tumor regions.

***SpaceFlow: Spatially Regularized Clustering***

SpaceFlow performs unsupervised spatial clustering using graph convolutional networks that incorporate spatial adjacency information [S2]. The method constructs a spatial graph wherein Visium spots are represented as nodes connected by edges based on physical proximity on the tissue section. The graph convolutional network learns low-dimensional embeddings that simultaneously capture transcriptomic similarity and spatial proximity, producing clusters that respect tissue architecture. SpaceFlow was run with default hyperparameters, and the number of initial clusters was determined by the algorithm's internal optimization. Fine-grained clusters were subsequently consolidated into the three principal spatial domains (Tumor, Boundary, Stroma) guided by CancerFinder malignancy scores and histological annotation.

***cell2location: Reference Atlas and Model Training***

The cell2location Bayesian deconvolution model was trained using a comprehensive single-cell RNA sequencing reference atlas compiled from published human liver and HCC datasets [S3]. The reference atlas encompassed T cells (including CD4+ and CD8+ subsets), B cells, natural killer cells, macrophages (including Kupffer cells and monocyte-derived subsets), dendritic cells, other myeloid lineages, hepatic stellate cells, cancer-associated fibroblasts, endothelial cells (tumor-associated and normal), fibroblasts, hepatocytes (normal and malignant), and cholangiocytes. The model was trained with default hyperparameters (4,000 training epochs for the reference model and 30,000 epochs for the spatial mapping model). Posterior estimates of absolute cell-type abundance per spot were extracted and normalized to proportions for cross-sample and cross-domain comparisons.

***PROGENy: Pathway Activity Estimation***

PROGENy (Pathway RespOnsive GENes) infers pathway activity from gene expression data based on consensus footprint gene signatures derived from large-scale perturbation experiments [S4]. Activity scores were computed for the following 14 signaling pathways: Androgen, EGFR, Estrogen, Hypoxia, JAK-STAT, MAPK, NF-kB, PI3K, p53, TGF-b, TNF-a, Trail, VEGF, and WNT. For each Visium spot, PROGENy scores were calculated using the top 500 most responsive genes per pathway as the footprint gene set.

***MISTy: Multiview Spatial Modeling***

MISTy (Multiview Intercellular SpaTial modeling framework) is an explainable multiview machine learning approach that decomposes spatially measured features into contributions from distinct spatial contexts [S5]. In our application, MISTy modeled pathway activity at each Visium spot as a function of cell-type abundances organized into two views: (1) the intrinsic view, representing cell-type proportions within the same spot, and (2) the paraview, representing cell-type proportions in the surrounding neighborhood. This decomposition enabled inference of whether specific cell populations drive signaling programs locally or from the broader tissue microenvironment.

***LIANA+: Ligand-Receptor Interaction Prioritization***

LIANA+ is a unified platform that integrates multiple ligand-receptor interaction databases (CellPhoneDB, CellChat, NATMI, Connectome, and others) with multiple scoring algorithms [S6]. By aggregating results across diverse resources and algorithmic approaches, LIANA+ generates consensus rankings of ligand-receptor pairs. We applied LIANA+ with its default aggregate scoring mode to prioritize interactions within and between the three spatial domains.

***stLearn: Spatially Resolved Cell-Cell Interaction Networks***

stLearn integrates gene expression data with physical inter-spot distances to construct cell-cell interaction networks grounded in tissue architecture [S7]. Unlike purely expression-based methods, stLearn down-weights interactions between spatially distant cell populations and up-weights those between co-localized populations, generating directional communication networks that reflect plausible signaling ranges. Interaction networks were generated for top-ranked ligand-receptor pairs from LIANA+ to visualize the cellular topology of specific signaling axes.

***STopover: Topological Spatial Overlap Analysis***

STopover applies topological data analysis to assess the spatial overlap between ligand and receptor expression domains [S8]. For each ligand-receptor pair, STopover computes a Jaccard composite score quantifying the degree to which the spatial expression patterns of the ligand and its cognate receptor co-localize within a defined tissue region. We applied STopover to the Boundary domain; interactions were ranked by Jaccard composite score, with those exceeding 0.3 designated strongly border-enriched and lower-scoring interactions with spatially concentrated boundary overlap classified as boundary-associated.

**Supplementary Table S1.** Clinical and pathological characteristics of the study cohort

| **Patient** | **Age** | **Sex** | **Etiology** | **Background liver** | **Fibrosis** | **pT** | **AFP (ng/mL)** | **PIVKA-II (mAU/mL)** |
| --- | --- | --- | --- | --- | --- | --- | --- | --- |
| P1 | 65 | F | HBV | Chronic hepatitis | Periportal | 2 | 3,071.4 | 718 |
| P2 | 55 | M | HBV | Chronic hepatitis | Septal | 2 | 39.1 | 27 |
| P3 | 63 | M | HBV | Chronic hepatitis | Portal | 2 | 3.5 | 109 |
| P4 | 65 | F | HBV | Chronic hepatitis | Periportal | 2 | 5.6 | 49 |
| P5 | 62 | M | HBV | Macronodular cirrhosis | — | 2 | 11,515.6 | 489 |
| P6 | 41 | M | HBV | Chronic hepatitis | Septal | 2 | 806.3 | 129 |
| P7 | 73 | F | HBV | Macro-/micronodular cirrhosis | — | 2 | 14.4 | 106 |
| P8 | 64 | F | NBNC | Chronic hepatitis | Septal | 1a | 2.0 | 25 |
| P9 | 70 | M | NBNC | Macro-/micronodular cirrhosis | — | 2 | 2.3 | 1,452 |
| P10 | 65 | M | NBNC | Moderate steatosis | Bridging | 2 | 3.8 | 15 |
| P11 | 70 | M | NBNC | Reactive hepatitis | Portal | 3 | 11.3 | 140 |

**Abbreviations:** AFP, alpha-fetoprotein; HBV, hepatitis B virus; HBV-LC, HBV-related liver cirrhosis; NBNC, non-B non-C; PIVKA-II, protein induced by vitamin K absence or antagonist-II; pT, pathological tumor stage.

**Supplementary Figure Legends**

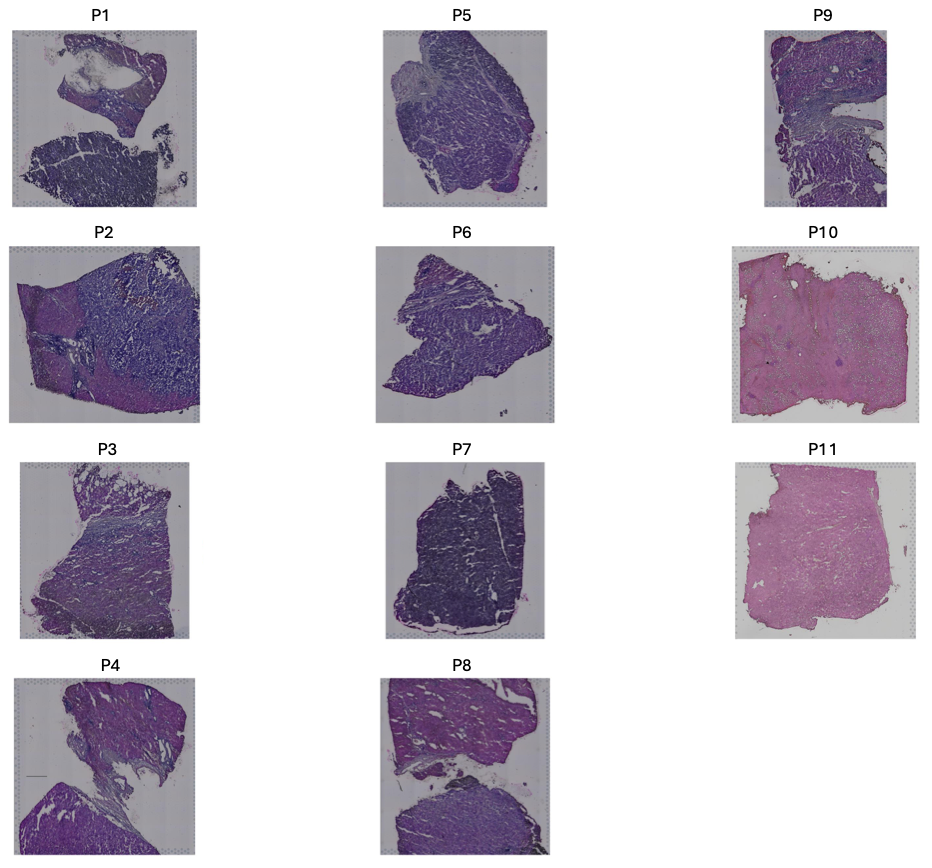

**Supplementary Fig. 1.** H&E-stained sections for all HCC specimens. Hematoxylin and eosin-stained tissue sections are shown for all 11 HCC specimens included in the spatial transcriptomics cohort. Specimens are labeled according to the final analysis identifiers P1 through P11. These images provide histologic context for the spatial domain annotation and downstream tumor-boundary-stroma analyses.

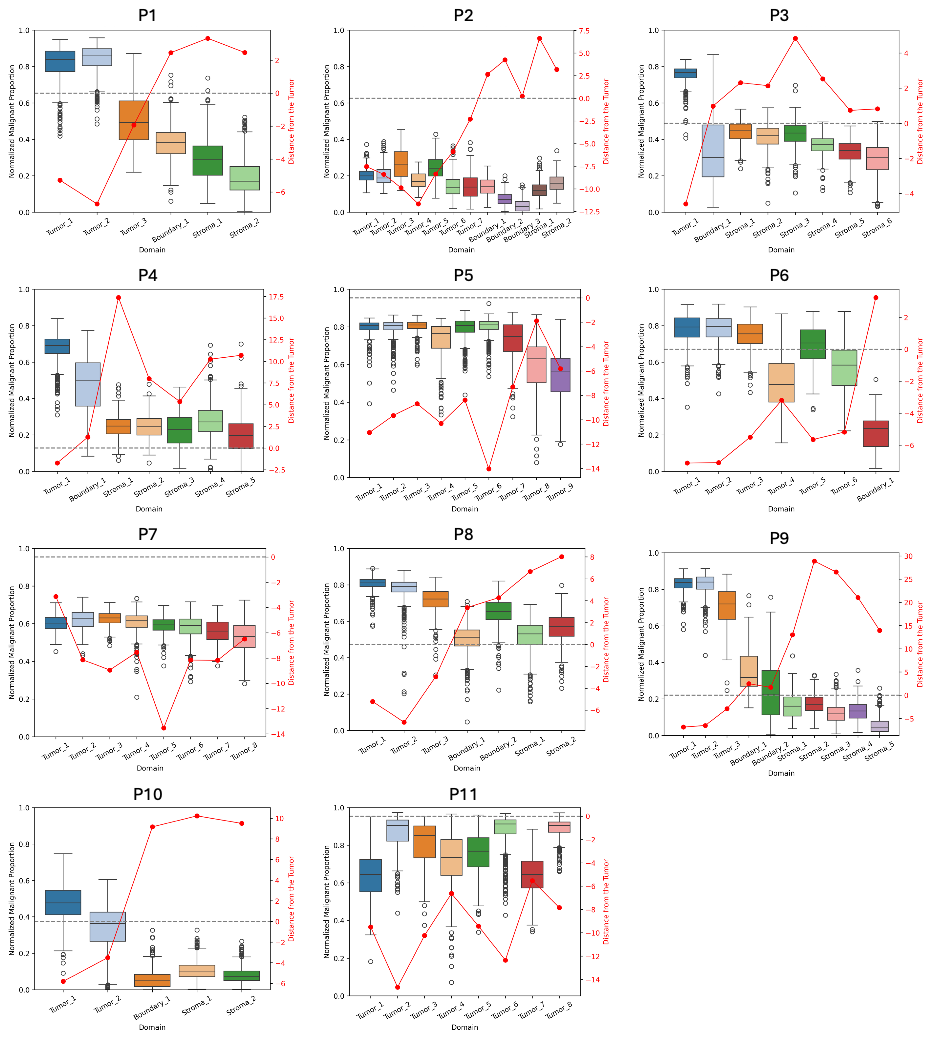

**Supplementary Fig. 2.** Validation of the tumor-border distance axis across all specimens. Boxplots of estimated malignant cell proportion are shown across ordered spatial subdomains for each of the 11 specimens. The red line overlay indicates the mean distance from the tumor for each subdomain. Across specimens, malignant cell proportions generally decreased with increasing distance from the tumor core, supporting the biological validity of the tumor-boundary-stroma spatial framework.

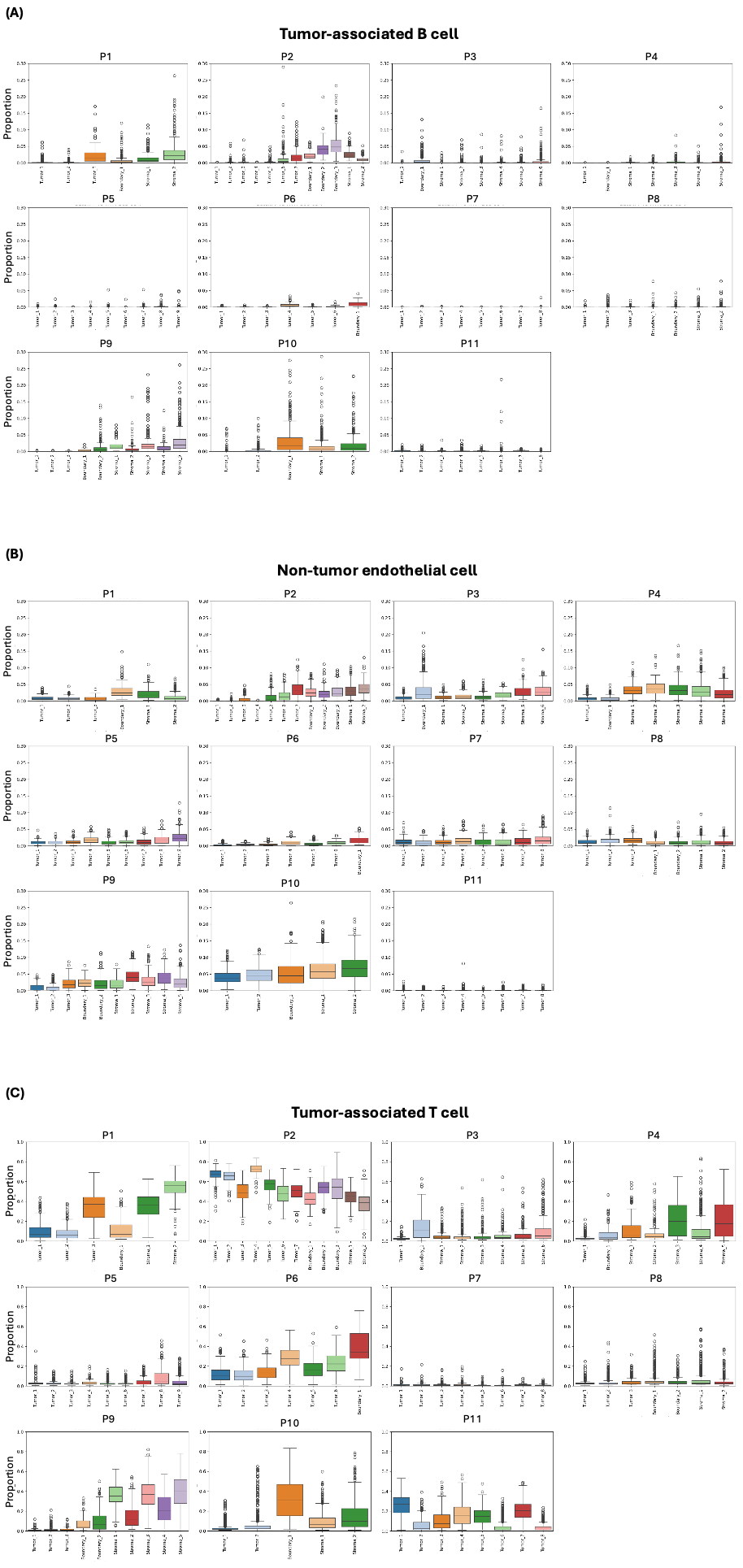

**Supplementary Fig. 3. Domain-level distributions of selected lymphoid and endothelial populations across all specimens.**

Boxplots of cell2location-inferred tumor-associated B cells (B_tumor), non-tumor endothelial cells (Endothelial_Normal), and tumor-associated T cells (T_tumor) are shown across Tumor, Boundary, and Stroma domains for all 11 specimens. Tumor-associated B and T cell labels refer to immune cell states in the reference atlas and should not be interpreted as malignant cells. Non-tumor endothelial cells are shown separately from tumor endothelial cells (TECs), which are presented in **Fig. 2**. These populations displayed more heterogeneous spatial distributions than the consistently boundary-enriched stromal and myeloid populations highlighted in **Fig. 2**.

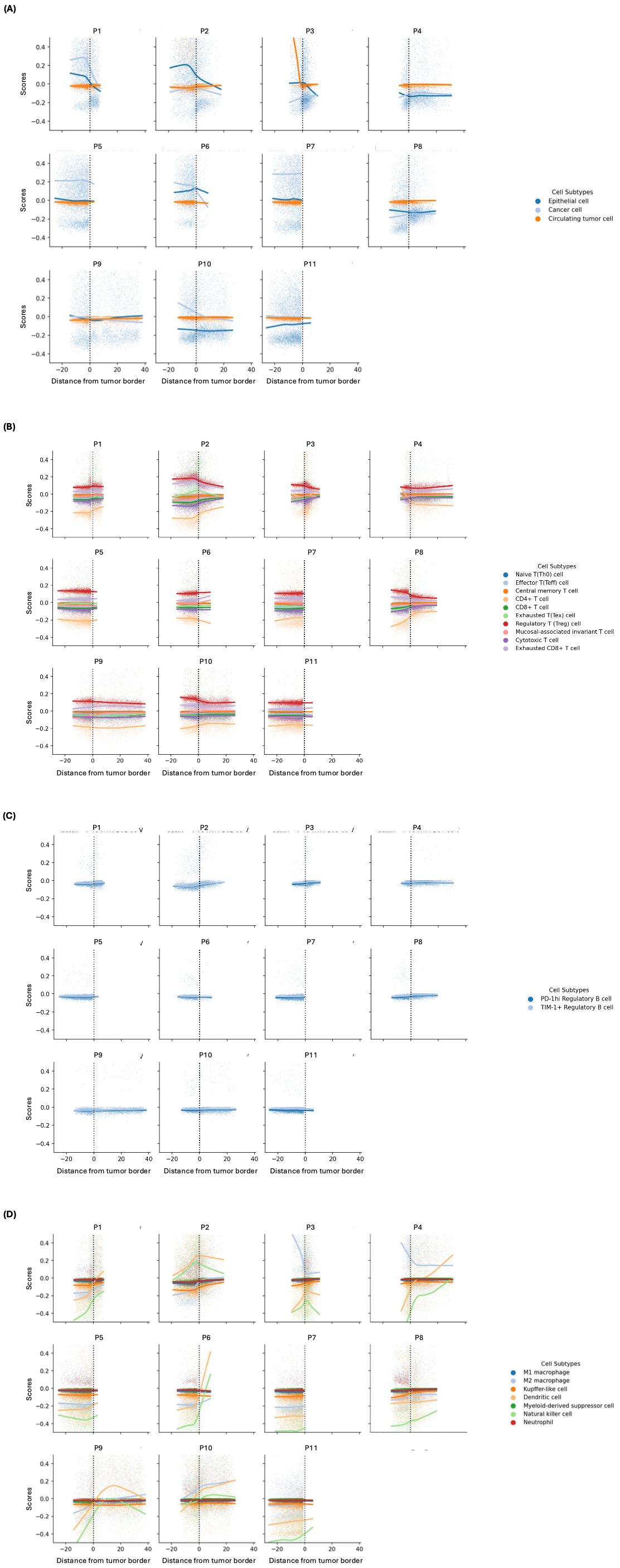

**Supplementary Fig. 4. Distance-to-border gradients of additional cell subtypes across all specimens.**

LOWESS trend curves are shown along the signed distance-to-border axis for each of the 11 specimens. The vertical dashed line at x = 0 marks the tumor border; negative values indicate tumor-side positions and positive values indicate stromal-side positions. a Gradients of epithelial, cancer, and circulating tumor cell-related populations. b Gradients of T cell subtypes. c Gradients of B cell subtypes. d Gradients of myeloid, macrophage, natural killer cell, and other immune-related populations. These analyses illustrate inter-patient heterogeneity in additional cell-state gradients beyond the core CAF, TAM, and TEC boundary-enriched populations highlighted in **Fig. 2**.

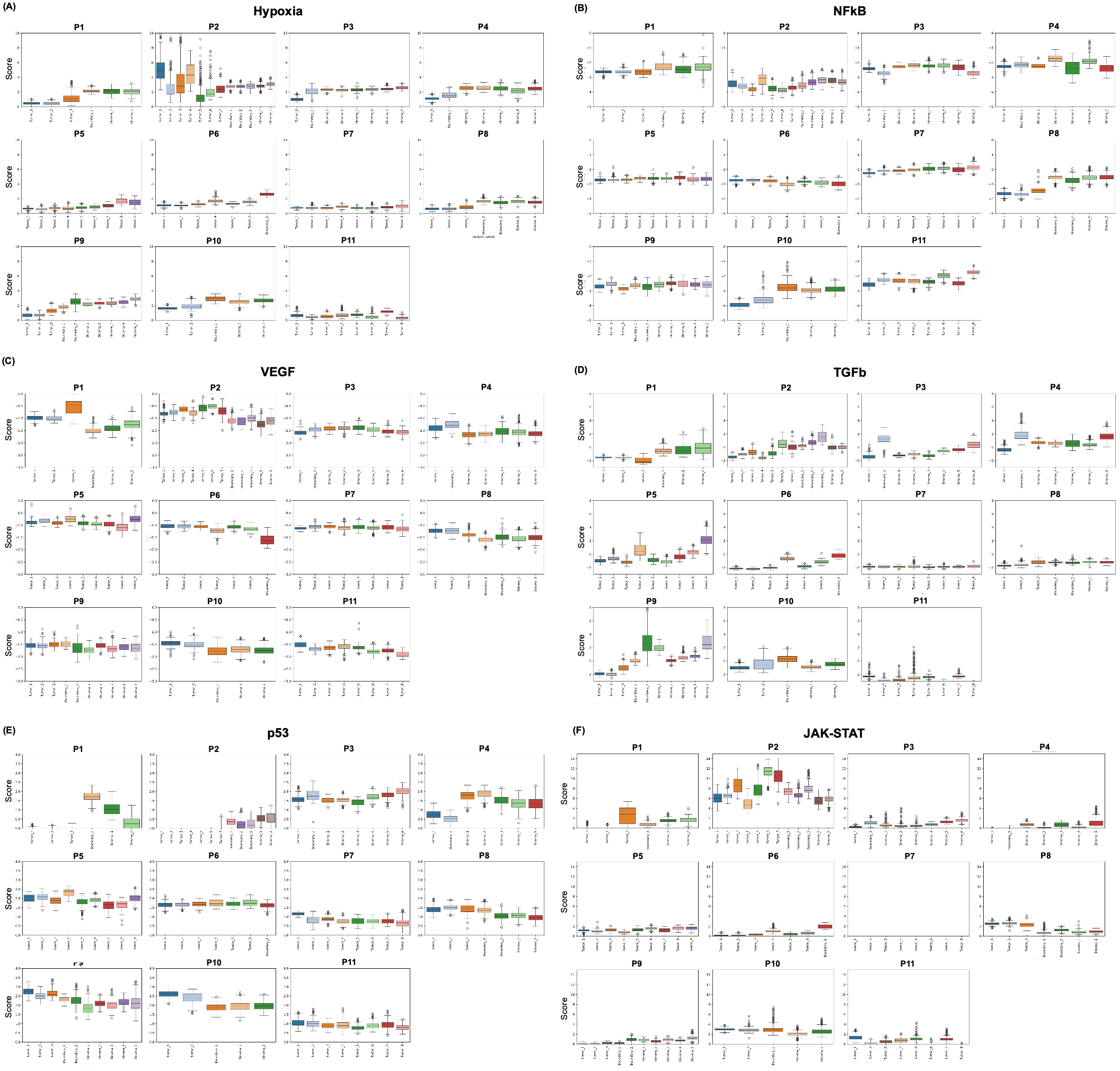

**Supplementary Fig. 5. Pathway activity distributions across spatial domains for all specimens.**

Boxplots of PROGENy-inferred pathway activity scores are shown across Tumor, Boundary, and Stroma domains for each of the 11 specimens. a Hypoxia, b NF-kB, c VEGF, d TGF-b, e p53, and f JAK-STAT pathway activity. These data complement the representative distance-to-border trend curves in **Fig. 3** by providing domain-level pathway summaries across the full cohort. JAK-STAT is included as an exploratory additional cytokine-associated pathway summary.
